# Predicted Coronavirus Nsp5 Protease Cleavage Sites in the Human Proteome: A Resource for SARS-CoV-2 Research

**DOI:** 10.1101/2021.06.08.447224

**Authors:** Benjamin M. Scott, Vincent Lacasse, Ditte G. Blom, Peter D. Tonner, Nikolaj S. Blom

## Abstract

**Background:** The coronavirus nonstructural protein 5 (Nsp5) is a cysteine protease required for processing the viral polyprotein and is therefore crucial for viral replication. Nsp5 from several coronaviruses have also been found to cleave host proteins, disrupting molecular pathways involved in innate immunity. Nsp5 from the recently emerged SARS-CoV-2 virus interacts with and can cleave human proteins, which may be relevant to the pathogenesis of COVID-19. Based on the continuing global pandemic, and emerging understanding of coronavirus Nsp5-human protein interactions, we set out to predict what human proteins are cleaved by the coronavirus Nsp5 protease using a bioinformatics approach.

**Results:** Using a previously developed neural network trained on coronavirus Nsp5 cleavage sites (NetCorona), we made predictions of Nsp5 cleavage sites in all human proteins. Structures of human proteins in the Protein Data Bank containing a predicted Nsp5 cleavage site were then examined, generating a list of 92 human proteins with a highly predicted and accessible cleavage site. Of those, 48 are expected to be found in the same cellular compartment as Nsp5. Analysis of this targeted list of proteins revealed molecular pathways susceptible to Nsp5 cleavage and therefore relevant to coronavirus infection, including pathways involved in mRNA processing, cytokine response, cytoskeleton organization, and apoptosis.

**Conclusions:** This study combines predictions of Nsp5 cleavage sites in human proteins with protein structure information and protein network analysis. We predicted cleavage sites in proteins recently shown to be cleaved *in vitro* by SARS-CoV-2 Nsp5, and we discuss how other potentially cleaved proteins may be relevant to coronavirus mediated immune dysregulation. The data presented here will assist in the design of more targeted experiments, to determine the role of coronavirus Nsp5 cleavage of host proteins, which is relevant to understanding the molecular pathology of SARS-CoV-2 infection.

## Background

Coronaviruses are major human and livestock pathogens, and are the current focus of international attention due to an ongoing global pandemic caused by severe acute respiratory syndrome coronavirus 2 (SARS-CoV-2). This recently emerged coronavirus likely originated in bats in China, before passing to humans in late 2019 through a secondary animal vector [1, 2]. Although 79% identical at the nucleotide level to SARS-CoV [2], differences in the infectious period and community spread of SARS-CoV-2 has caused a greater number of cases and deaths worldwide [3, 4]. Individuals infected by SARS-CoV-2 can develop COVID-19 disease which primarily affects the lungs, but can also cause kidney damage, coagulopathy, liver damage, and neuropathy [5–10]. Hyperinflammation, resulting from dysregulation of the immune response to SARS-CoV-2 infection, has emerged as a leading hypothesis regarding severe COVID-19 cases, which may also explain the diverse and systemic symptoms observed [11–14].

Similar to other coronaviruses, once a cell is infected, the 5’ portion of the SARS-CoV-2 (+)ssRNA genome is translated into nonstructural proteins (Nsps) required for viral replication, which are expressed covalently linked to one another (Fig. 1a) [15]. This polyprotein must therefore be cleaved to free the individual Nsps, which is performed by two virally encoded proteases: Nsp3/papain-like protease (PLpro) and Nsp5/Main Protease (Mpro)/3C-like protease (3CLpro). Nsp5 is responsible for the majority of polyprotein cleavages and its function is conserved across coronaviruses [16, 17], making it a key drug target as its inhibition impedes viral replication (reviewed by [18]).

**Fig. 1.**
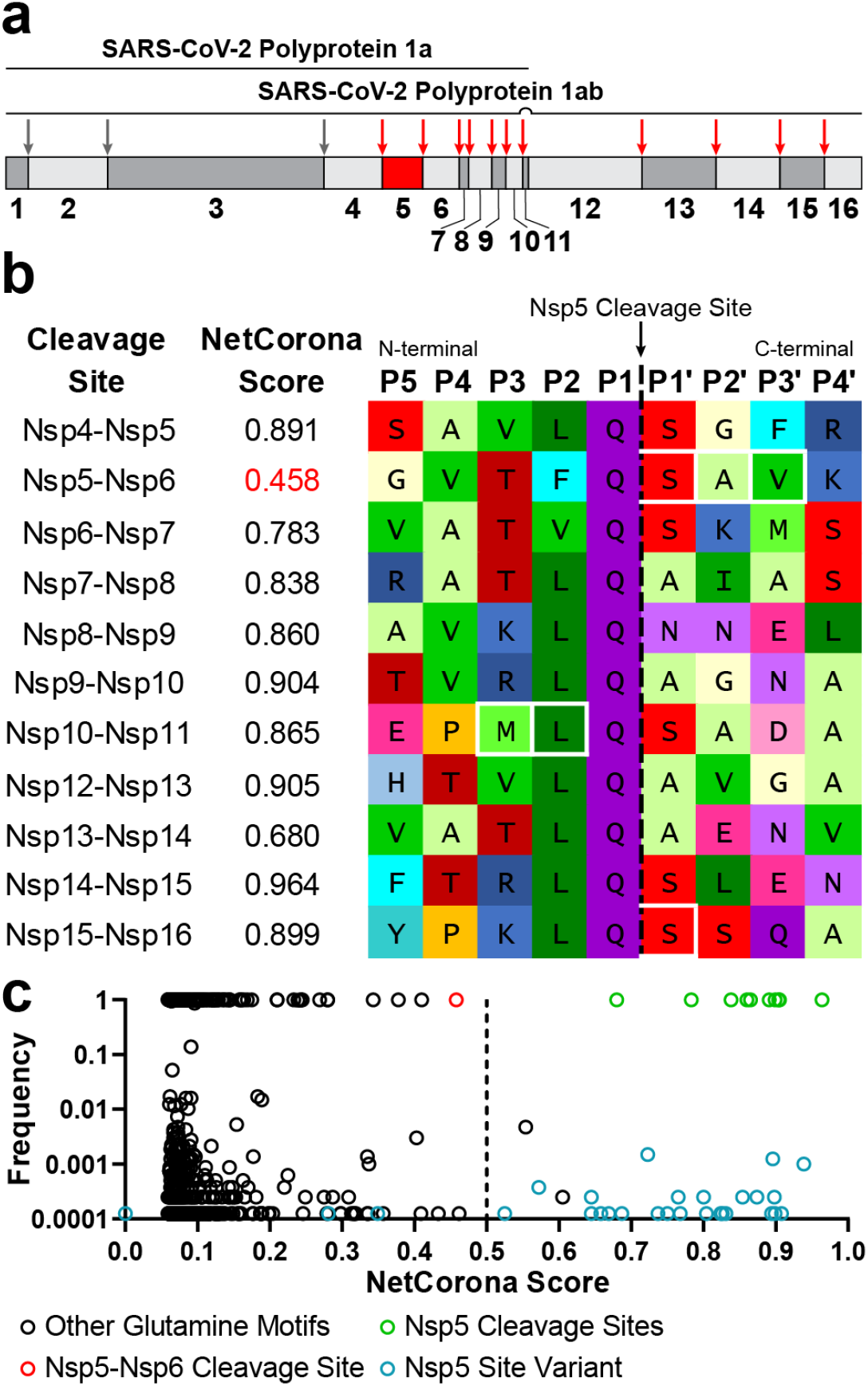
**(a) The SARS-CoV-2 1a and 1ab polyproteins**. 1a contains Nsp1-Nsp11, 1ab contains Nsp1-Nsp16 with Nsp11 skipped by a −1 ribosomal frameshift. Nsp5 and its cleavage sites are indicated with red arrows. Nsp3 cleavage sites are indicated with grey arrows. **(b) SARS-CoV-2 native Nsp5 cleavage motifs.** NetCorona scores are indicated, and residues in white boxes differ from SARS-CoV. **(c) SARS-CoV-2 1ab sequences scored with NetCorona.** Scores and frequency were determined for all P5-P4’ motifs surrounding glutamine residues in 8017 patient-derived SARS-CoV-2 sequences. Known Nsp5 cleavage sites are indicated in green, while mutations at a Nsp5 cleavage site are indicated in blue. The Nsp5-Nsp6 cleavage site is indicated in red, and all other glutamine motifs are indicated in black.

All coronavirus Nsp5 proteases identified to date are cysteine proteases in the chymotrypsin family, which primarily cleave peptides at P2-P1-P1’ residues leucine-glutamine-alanine/serine [16, 17, 19, 20], where the cleavage occurs between the P1 and P1’ residues. Nsp5 forms a homodimer for optimal catalytic function but may function as a monomer when processing its own excision from the polyprotein [21–23]. SARS-CoV-2 Nsp5 shares 96.1% sequence identity with SARS-CoV Nsp5 and has similar substrate specificity *in vitro*, but SARS-CoV-2 Nsp5 accommodates more diverse residues at substrate position P2 and may have a higher catalytic efficiency [22–25].

Coronavirus proteases also manipulate the cellular environment of infected cells to favor viral replication [26, 27], and disrupt host interferon (IFN) signaling pathways to suppress the anti-viral response of the innate immune system (reviewed by [28–30]). The role of coronavirus protease Nsp3 as an IFN antagonist has been well documented, including SARS-CoV-2 Nsp3 [31, 32]. Although Nsp3 proteolytic activity contributes to IFN antagonism, it is the deubiquitinating and deISGylating activities of Nsp3 that are primarily responsible [32–39]. In contrast, fewer examples of Nsp5 mediated disruption of host molecular pathways have been identified, and all are a result of its proteolytic activity [40–43].

SARS-CoV-2 Nsp5 antagonism of IFN is not yet clear [31, 44, 45], but *in vitro* evidence supports SARS-CoV-2 Nsp5 mediated cleavage of TAB1 and NLRP12 which are involved in innate immunity [46]. Hundreds of potentially cleaved peptides containing the Nsp5 consensus sequence appeared when lysate from human cells were incubated with recombinant Nsp5 from SARS-CoV, SARS-CoV-2, or hCoV-NL63, indicating a significant potential for Nsp5 mediated disruption of host proteins [47]. Similarly, the abundance of potentially cleaved peptides containing a Nsp5 consensus sequence was increased in cells infected *in vitro* with SARS-CoV-2, which was dependent on the cell type studied [48]. Knock down or inhibition of some of these human proteins likely cleaved by Nsp5, suppressed SARS-CoV-2 replication *in vitro*, suggesting that targeted host protein proteolysis is involved in viral replication [48]. Many other SARS-CoV-2 Nsp5-host protein interactions have been identified using proximity labeling and co-immunoprecipitation [49–54], but it is unknown if these interactions lead to Nsp5 mediated cleavage. Indeed, *in vitro* studies may miss Nsp5-host protein interactions due to cleavage of the host protein upon Nsp5 binding [49], and because individual cell types only express a limited set of human proteins. A proteome-wide prediction of coronavirus Nsp5 mediated cleavage of human proteins is therefore relevant to understanding COVID-19 pathogenesis, and how coronaviruses in general disrupt host biology.

The neural network NetCorona was previously developed in 2004, and was trained on a dataset of Nsp5 cleavage sites from seven coronaviruses including SARS-CoV [55]. NetCorona outperforms traditional consensus motif-based approaches for identifying cleavage sites, and based on the similar specificities of SARS-CoV and SARS-CoV-2 Nsp5, we believed it could be applied to the study of SARS-CoV-2 Nsp5 interactions with human proteins. However, NetCorona only analyzes the primary amino acid sequence to predict cleavage sites, which lacks information about the 3D structure of the folded protein, and therefore how exposed a predicted cleavage site is to a protease. In particular, the solvent accessibility of a peptide motif is closely related to proteolytic susceptibility [56, 57], and *in silico* measurement of solvent accessibility has previously been used to help predict proteolysis [58–60].

In this study we used NetCorona to make predictions of Nsp5 cleavage sites across the entire human proteome, and additionally analyzed available protein structures *in silico* to identify highly predicted cleavage sites. We extended this analysis to examine subcellular and tissue expression patterns of the proteins predicted to be cleaved, and applied protein network analysis to identify potential key pathways disrupted by Nsp5 cleavage. Predicted Nsp5 cleavage sites in human proteins were similar to those recently identified *in vitro*, and human proteins predicted to be cleaved by Nsp5 were found to be involved in molecular pathways that may be relevant to the pathogenesis of COVID-19 disease.

## Results

### Evaluating NetCorona Performance with the SARS-CoV-2 Polyprotein

As the NetCorona neural network was not trained on the SARS-CoV-2 polyprotein sequence (Fig. 1a), we first examined if the 11 polyprotein cleavage sites homologous to SARS-CoV would be correctly scored as cleaved (NetCorona score >0.5). Due to the high sequence similarity with SARS-CoV, there were only three cleavage sites containing different residues (Fig. 1b). The mean NetCorona score for 10 out of the 11 SARS-CoV-2 Nsp5 cleavage sites was 0.859 (SD = 0.08), indicating highly predicted cleavages. The cleavage site at Nsp5-Nsp6 was classified as uncleaved, with a score of 0.458. SARS-CoV contains the same unique phenylalanine at position P2 of Nsp5-Nsp6, but with different P1’-P3’ residues, and received a marginal score of 0.607 in the original NetCorona paper [55]. Phenylalanine at P2 is not found in other coronaviruses that infect humans [20, 61], nor in the other viruses used to train NetCorona, which contributed to these low scores. A P2 phenylalanine may be intentionally unfavorable at the Nsp5-Nsp6 cleavage site, to assist in its autoprocessing from the polypeptide, by limiting the ability of the cleaved peptide’s C-terminus to bind the Nsp5 active site [61]. Interestingly, the swapped identity of the P3 and P2 residues at the SARS-CoV-2 Nsp10-Nsp11 cleavage site resulted in a higher score versus SARS-CoV (0.865 vs 0.65), due to leucine being more common at P2 versus methionine. This mutation may result in a more rapid cleavage at this site in SARS-CoV-2 versus SARS-CoV, as Nsp5 favors leucine above all other residues at P2 [17, 25].

To investigate if NetCorona can distinguish between cleaved and uncleaved motifs, NetCorona scores for all glutamine motifs in the SARS-CoV-2 1ab polyprotein were also determined. To gather context from the ongoing pandemic and to investigate glutamine motifs across different viral variants, 8017 SARS-CoV-2 1ab polyprotein sequences obtained from patient samples were scored with NetCorona (Fig. 1c, Additional File 1: Table S1). Apart from two motifs present in only 40 sequences, all glutamine motifs not naturally processed by Nsp5 received a NetCorona score <0.5, indicating they were correctly predicted not to be cleaved. Mutations at native Nsp5 cleavage sites were also rare, with only 28 such mutated cleavage sites present in 63 sequences. Except for three mutations present in one sequence each, mutations at native Nsp5 cleavage sites were conservative and only modestly changed the NetCorona score. One sequence contained a histidine at Nsp8-Nsp9 P1 (QIO04366), resulting in NetCorona not scoring the motif. SARS-CoV and SARS-CoV-2 Nsp5 may be able to cleave motifs with histidine at P1, albeit with reduced efficiency [17, 47].

These combined results indicate that despite NetCorona not being trained on the SARS-CoV-2 sequence, it was able to correctly distinguish between cleaved versus uncleaved motifs in the 1ab polyprotein, except for Nsp5-Nsp6. The rarity of mutated canonical cleavage sites and mutations introducing new cleavage sites (0.8% and 0.5% of sequences respectively), indicates stabilizing selection for a distinction between Nsp5 cleavage sites and all other glutamine motifs.

### NetCorona Predictions of Nsp5 Cleavage Sites in the Human Proteome

To generate a global view of Nsp5 cleavage sites in the human proteome, datasets were batch analyzed using NetCorona (Fig. 2). Every 9-residue motif flanking a glutamine was scored, where glutamine acts as P1 and four resides were analyzed on either side (P5-P4’). Using a NetCorona score cutoff of >0.5, 15057 proteins (~20%) in the “All Human Proteins” dataset contained a predicted cleavage site, 6056 (~29%) proteins in the “One Protein Per Gene”, and 2167 (~32%) proteins in the “Proteins With PDB” dataset (Additional File 1: Table S2-S4, raw data sets in Additional File 2-4).

**Fig. 2.**
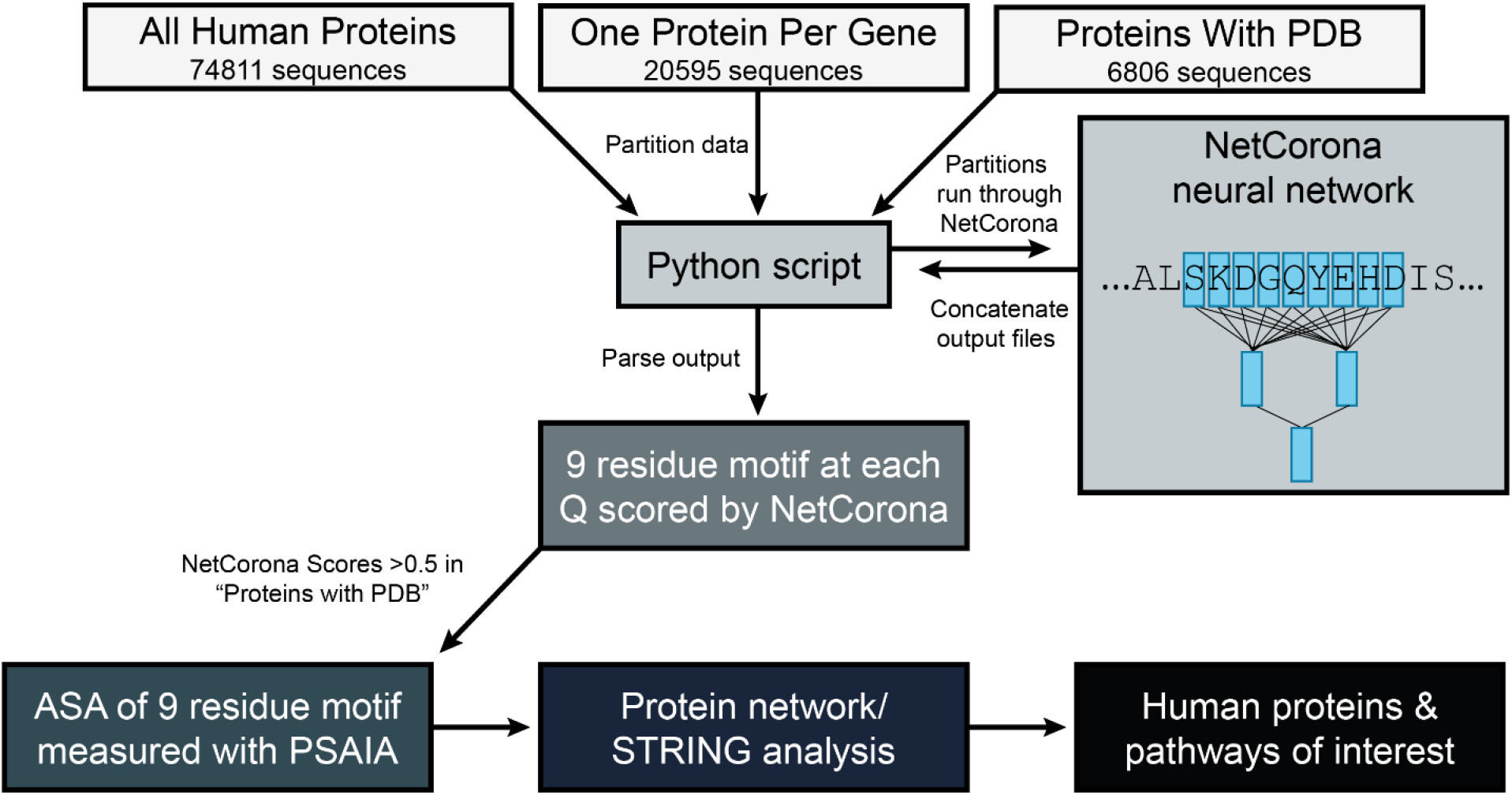
Overview of approach to predicting Nsp5 cleavage sites in human proteins. Three datasets of human protein sequences were analyzed by the NetCorona neural network. NetCorona assigned scores (0 – 1.0) to the 9 amino acid motif surrounding every glutamine residue in the datasets, where a score >0.5 was inferred to be a possible cleavage site. PDB files associated with predicted cleaved proteins were analyzed using the Protein Structure and Interaction Analyzer (PSAIA) tool, which output the accessible surface area (ASA) of each predicted 9 amino acid cleavage motif. Proteins with highly predicted Nsp5 cleavage sites were then analyzed using STRING, which provided information on tissue expression, subcellular localization, and performed protein network analysis. Human proteins and molecular pathways of interest containing a predicted Nsp5 cleavage site were then flagged for potential physiological relevance.

To help interpret these results, we compared the output from “One Protein Per Gene” to proteins that have been directly tested *in vitro* for cleavage by a coronavirus Nsp5 protease (Additional File 1: Table S5). There are six human proteins where cleavage sites have been mapped to the protein sequence (GOLGA3, NEMO, NLRP12, PAICS, PNN, TAB1) [40, 46, 48], and also two proteins from pigs (NEMO, STAT2) [41, 43], and one from cats (NEMO) [42]. NetCorona accurately scored 6 out of the 12 unique cleavage sites mapped in these proteins. NetCorona struggled with an identical cleavage motif at Q231 in NEMO from cats, pigs, and humans, which contains an uncommon valine at P1’. Interestingly, NetCorona predicted a cleavage site in PNN at Q495, which was not identified in the original study but matches the size of a reported secondary cleavage product [48].

Instances where NetCorona predicted cleavages but they are not observed *in vitro* are also relevant to interpreting the full proteome results. NetCorona predicted cleavage sites in 22 of the 71 proteins Moustaqil *et al*. studied, however only TAB1 and NLRP12 were observed to be cleaved by SARS-CoV-2 Nsp5 [46]. NetCorona predicted three cleavage sites in TAB1 and two in NLRP12, but just one predicted site in each protein matched the mapped cleavage sites.

Many other potential cleavage sites have been identified by Koudelka *et al*. [47], where N-terminomics was used to identify possible cleavage sites, after cell lysate was incubated with various coronavirus Nsp5 proteases. Out of the 383 unique peptides where a glutamine was at P1, NetCorona predicted that 167 (44%) of them would be cleaved (Additional File 1: Table S6). Meyer *et al*. similarly used N-terminomics to study potential Nsp5 cleavage events, following *in vitro* infection with SARS-CoV-2 [48]. They identified 12 motifs in human proteins that were likely cleaved by Nsp5, of which NetCorona predicted 8 of these to be cleaved (Additional File 1: Table S7).

Several SARS-CoV-2 human protein interactomes have been made available [49–54], where interactions between Nsp5 and human proteins have been reported. Interactions identified by Samavarchi-Therani *et al*. were the most numerous, and the data was well annotated [51], which enabled a comparison to our results. These interaction scores, which varied depending on where the BioID tag was located on Nsp5 (Nsp5 C-term, N-term, or N-term on the C145A catalytically inactive mutant), were plotted against the NetCorona score from our study, which is illustrated in Additional File 5: Figure S1 (raw data in Additional File 1: Table S8). Although statistically significant, the negative correlation between the strength of the Nsp5-human protein interaction and the maximum NetCorona score was small: ρ ranged from −0.18 to −0.29, r^2^ ranged from 0.03 to 0.08, depending on where the BioID tag was located on Nsp5. When examining only the human proteins with a positive interaction score, the mean NetCorona score ranged from 0.35 to 0.38 (SD = 0.25). Thus, Nsp5-human protein interactions identified *in vitro* by Samavarchi-Therani *et al*. did not reflect an increased likelihood of cleavage predicted by NetCorona.

### Structural Characterization of Predicted Nsp5 Cleavage Sites

We next sought to incorporate available structural information of potential protein substrates into our analysis, to address the discrepancy between the cleavage events predicted by NetCorona, and mapped cleavage sites that have been directly observed *in vitro*. The “Proteins With PDB” dataset contains only human proteins that have a solved structure available in the RCSB Protein Data Bank (PDB), however technical limitations for solving protein structures means that certain protein domains, such as transmembrane and disordered regions, may be underrepresented [62]. To investigate if the available PDB structures contained a biased distribution of NetCorona scores, similarity between the distribution of NetCorona scores for “Proteins With PDB” and proteins in the other two datasets was assessed through the non-parametric KS test (Fig. 3a). There was insufficient evidence to reject the null hypothesis that the distribution of scores for “Proteins With PDB” proteins was equivalent to scores for “All Human Proteins” and “One Protein Per Gene” (p=0.121 and p=0.856, respectively), indicating that there was not significant bias in the distribution of NetCorona scores.

**Fig. 3.**
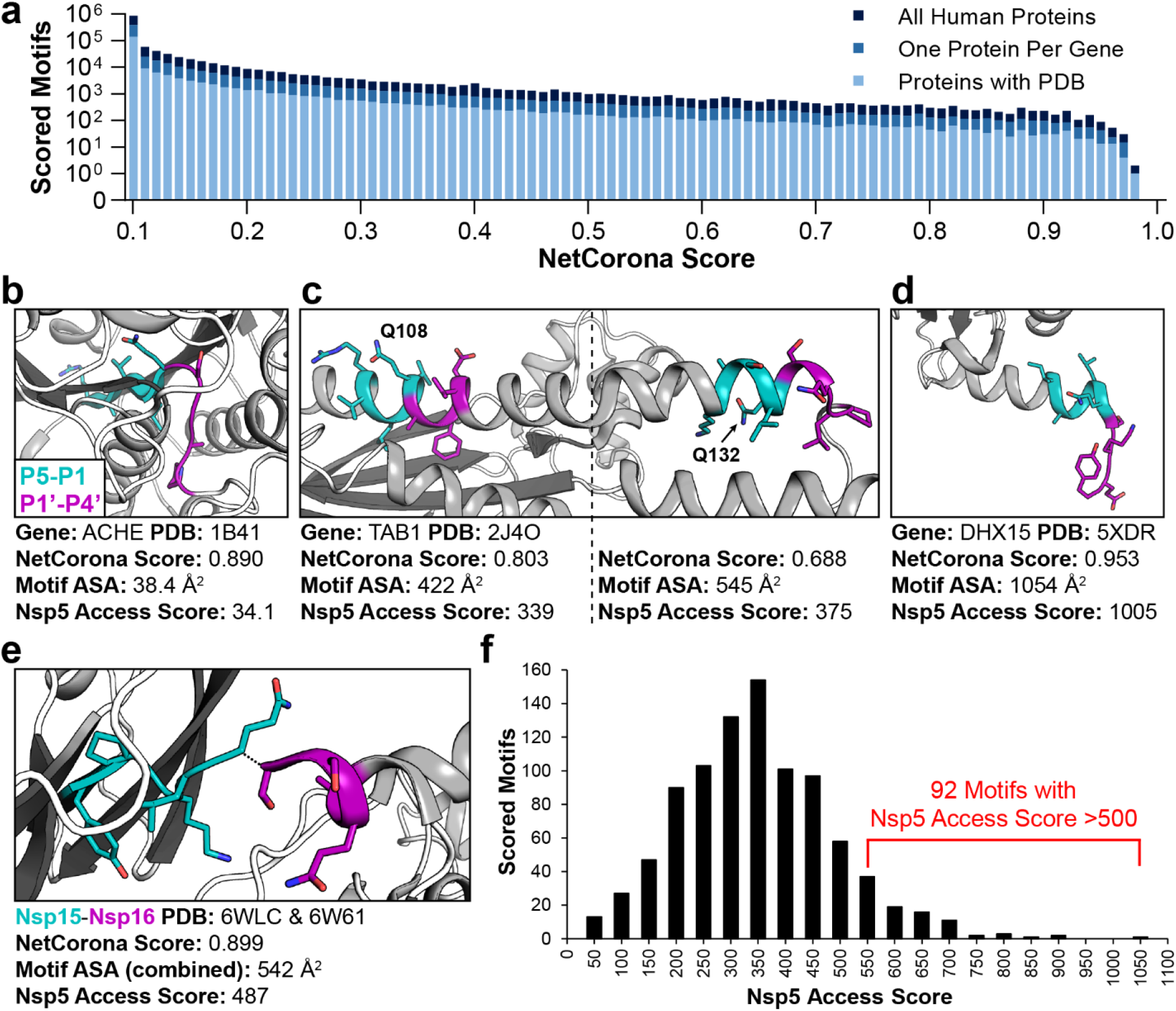
Structural analysis of predicted and known Nsp5 cleavage motifs. **(a)** NetCorona scores are shown for all P5-P4’ motifs surrounding glutamine residues in three datasets of human proteins, binned by score differences of 0.01. The distributions of scores were not statistically different from one another. **(b)** Despite a high NetCorona score in ACHE, the motif’s location in the core of the protein leads to a low Nsp5 access score. **(c)** TAB1 contains several motifs predicted to be cleaved, including at Q108 and Q132. The Nsp5 access score is slightly higher for the Q132 motif due to the greater accessible surface area (ASA). **(d)** DHX15 contains the motif with the highest Nsp5 access score observed in the human proteins studied, located on the C-terminus of the protein. **(e)** SARS-CoV-2 proteins Nsp15 and Nsp16 contain the native Nsp5 cleavage motif with the lowest Nsp5 access score calculated (487), which helped provide a cut-off to Nsp5 access scores in human proteins. **(f)** The Nsp5 access score of human protein motifs are indicated, binned by score differences of 50. 92 motifs in 92 unique human proteins have a Nsp5 access score >500.

NetCorona scores are derived from the primary amino acid sequence, but targeted proteolysis is also dependent on the 3D structural context of the potential substrate peptide within a protein [56, 57]. Many methods have been developed to quantify this structural context *in silico*, and solvent accessibility has been shown to be a strong predictor of proteolysis [57]. Accessible surface area (ASA) is commonly used to measure solvent accessibility, where a probe that approximates a water molecule is rolled around the surface of the protein, and the path traced out is the accessible surface [63]. Thin slices are then cut through this path, to calculate the accessible surface of individual atoms. After obtaining PDB files containing motifs predicted to be cleaved by NetCorona, the total ASA of each 9 amino acid motif was calculated using Protein Structure and Interaction Analyzer (PSAIA) [64]. This ASA was then multiplied by the motif’s NetCorona score to provide a “Nsp5 access score”, which represents both the solvent accessibility and substrate sequence preference. A Nsp5 access score was obtained for 914 glutamine motifs in 794 unique human proteins (Additional File 1: Table S9), with the process for selecting PDB files to analyze listed in Additional File 6.

Specific examples are presented to illustrate the utility of the Nsp5 access score (Fig. 3b-e). Acetylcholinesterase (ACHE) contains a motif at Q259 that was highly scored by NetCorona (0.890), but due to its presence in a tightly packed beta sheet in the core of the protein, the low ASA (38.4 Å^2^) results in a similarly low Nsp5 access score (34.1) and is therefore unlikely to be cleaved by Nsp5 (Fig. 3b). TGF-beta-activated kinase 1 (TAB1) is one of the few human proteins with a structure and experimental evidence of SARS-CoV-2 cleavage at specific sites (Q132 and Q444) [46]. As illustrated in Fig. 3c, the nearby motif at Q108 was scored higher than Q132 by NetCorona, but the greater ASA of the Q132 motif contributes to a higher Nsp5 access score, which matches the experimental evidence. The human protein with the highest Nsp5 access score was DEAH box protein 15 (DHX15), as the motif surrounding Q788 was both highly scored by NetCorona and its location proximal to the C-terminus of the protein makes it highly solvent exposed (Fig. 3d).

### Rationale for Nsp5 Access Score Cut-Off

To focus analysis on human proteins most likely to be cleaved by Nsp5, we determined a relevant cut-off to the Nsp5 access score. Using available structures and homology models, the Nsp5 access score of SARS-CoV-2 native cleavage sites was calculated, which ranged from 487 (Nsp15-Nsp16) to 923 (Nsp4-Nsp5) (Additional File 1: Table S10). The Nsp15-Nsp16 site (Fig. 3e) had a surprisingly low ASA (542 Å^2^) versus the other SARS-CoV-2 cleavage sites analyzed (mean of others 890 Å^2^, SD = 102 Å^2^), and as compared to the P5-P4’ residues of known substrates of other proteases in the chymotrypsin family (mean 678 Å^2^, SD = 297 Å^2^) (Additional File 1: Table S11).

As previously noted, NetCorona predicted cleavage sites in 22 of the 71 proteins Moustaqil *et al*. studied, but cleavages were only observed *in vitro* in two proteins [46]. Nsp5 access scores could be provided for 9 unique motifs from the 22 proteins NetCorona predicted to be cleaved, the mean of which was 332 (SD = 143). The sum of this mean and one standard deviation gives a Nsp5 access score of 475. 30 cleavage sites identified *in vitro* by Koudelka *et al*. could be assigned a Nsp5 access score [47], the mean of which was 381 (SD = 150). The sum of this mean and one standard deviation gives a Nsp5 access score of 531.

Based on these comparisons to available experimental data, a Nsp5 access score cut-off of 500 was selected, which is further illustrated in Additional File 7: Figure S2 (full data in Additional File 1: Table S12). This cut-off accommodates motifs with marginal NetCorona scores (~0.5) but maximally observed ASA (~1000 Å^2^), and the opposite scenario where a low ASA comparable to Nsp15-Nsp16 (~500 Å^2^) is matched with a high NetCorona score (~0.9). 92 motifs in 92 human proteins were found to have a Nsp5 access score >500 (Fig. 3f), which were forwarded to the next rounds of analysis.

### Analysis of Tissue Expression and Subcellular Localization of Predicted Cleaved Proteins

Proteins with a Nsp5 access score above 500 were imputed in STRING within the Cytoscape environment [65–67]. The STRING app computes protein network interaction by integrating information from publicly available databases, such as Reactome and Uniprot. Through textmining of the articles reported in those databases, it also compiles scores for multiple tissues and cellular compartment. The nucleus and cytosol were the top locations for human proteins with a highly predicted Nsp5 cleavage site (Fig. 4a), and the highest expression was in the nervous system and liver (Fig. 4b). The mean or summed expression score did not correlate with the Nsp5 access score (ρ = 0.03 and 0.05 respectively), nor was there a correlation between the Nsp5 access score and subcellular localization scores (ρ = −0.08 for mean and −0.17 for sum).

**Fig. 4.**
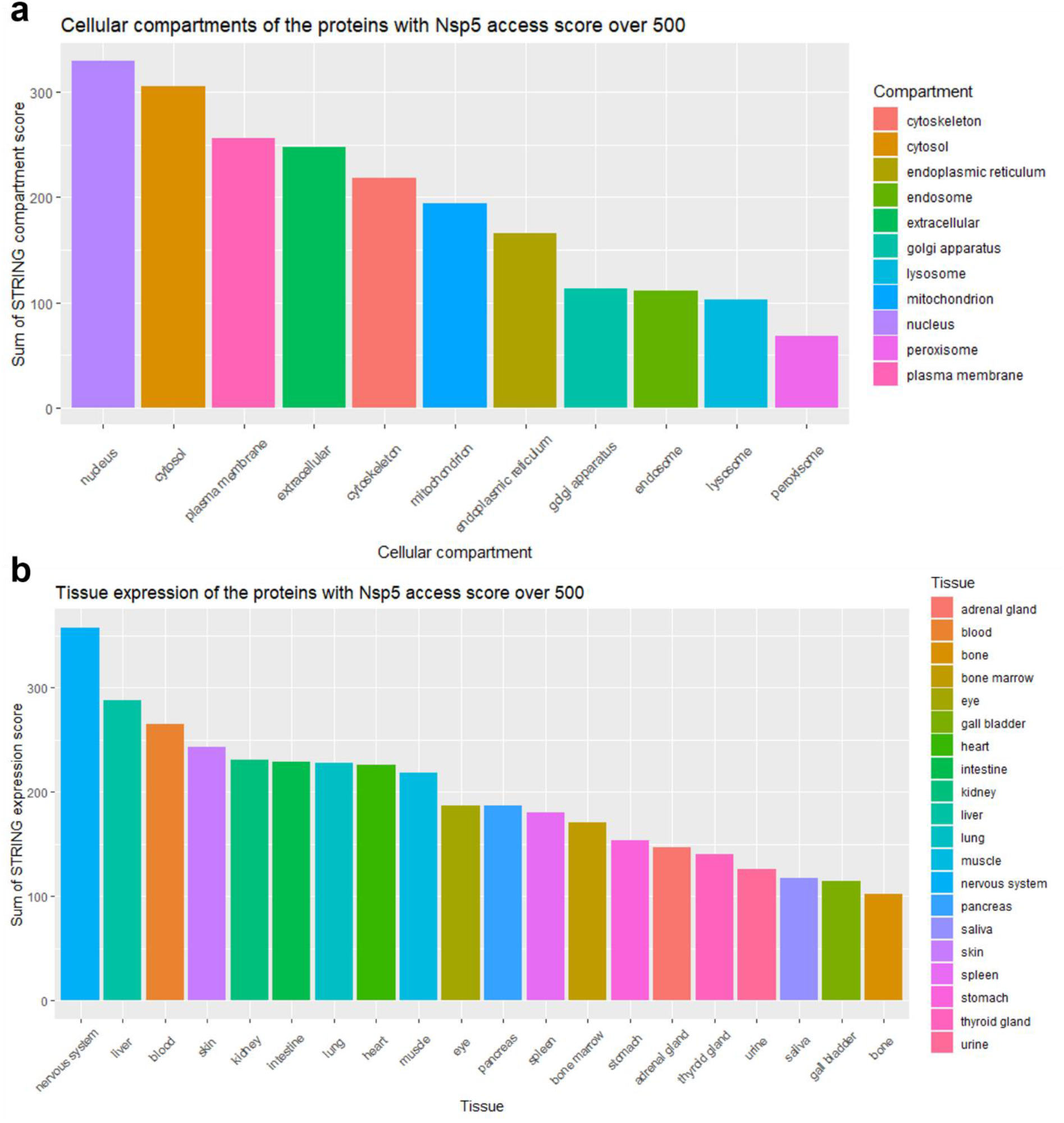
Sum of the compartment score **(a)** or expression score **(b)** of all human proteins with a Nsp5 access score above 500 (92 proteins). Both the compartment and the expression score were obtained from STRING based on text-mining and database searches.

Studies of the subcellular localization of coronavirus Nsps provide insight into where Nsp5 may exist in infected cells, and thus what human proteins it may be exposed to. Flanked by transmembrane proteins Nsp4 and Nsp6 in the polyprotein, Nsp5 is exposed to the cytosol when first expressed, where it colocalizes with Nsp3 once released [68–70]. Recent studies have indicated that SARS-CoV-2 Nsp5 activity can be detected throughout the cytosol of a patient’s cells *ex vivo* [25], and Nsp5 is also found in the nucleus and ER [51, 71].

Through the Human Protein Atlas (HPA), we obtained information on protein expression in tissue by immunohistochemistry (IHC) together with intracellular localization obtained by confocal imaging for most of the proteins in our dataset [72]. Proteins that are not found in the same cellular compartment as Nsp5, or where intracellular localization was unknown, were filtered out. Out of the initial 92 proteins with a Nsp5 access score over 500 and based on current knowledge, only 48 proteins were likely to be found in the same cellular compartment as Nsp5 (Fig. 5, Additional File 1: Table S13-14), indicating the greatest potential for interacting with and being cleaved by the protease. Proteins involved in apoptosis, such as CASP2, E2F1, and FNTA, had both a high Nsp5 access score and an above average expression.

**Fig. 5.**
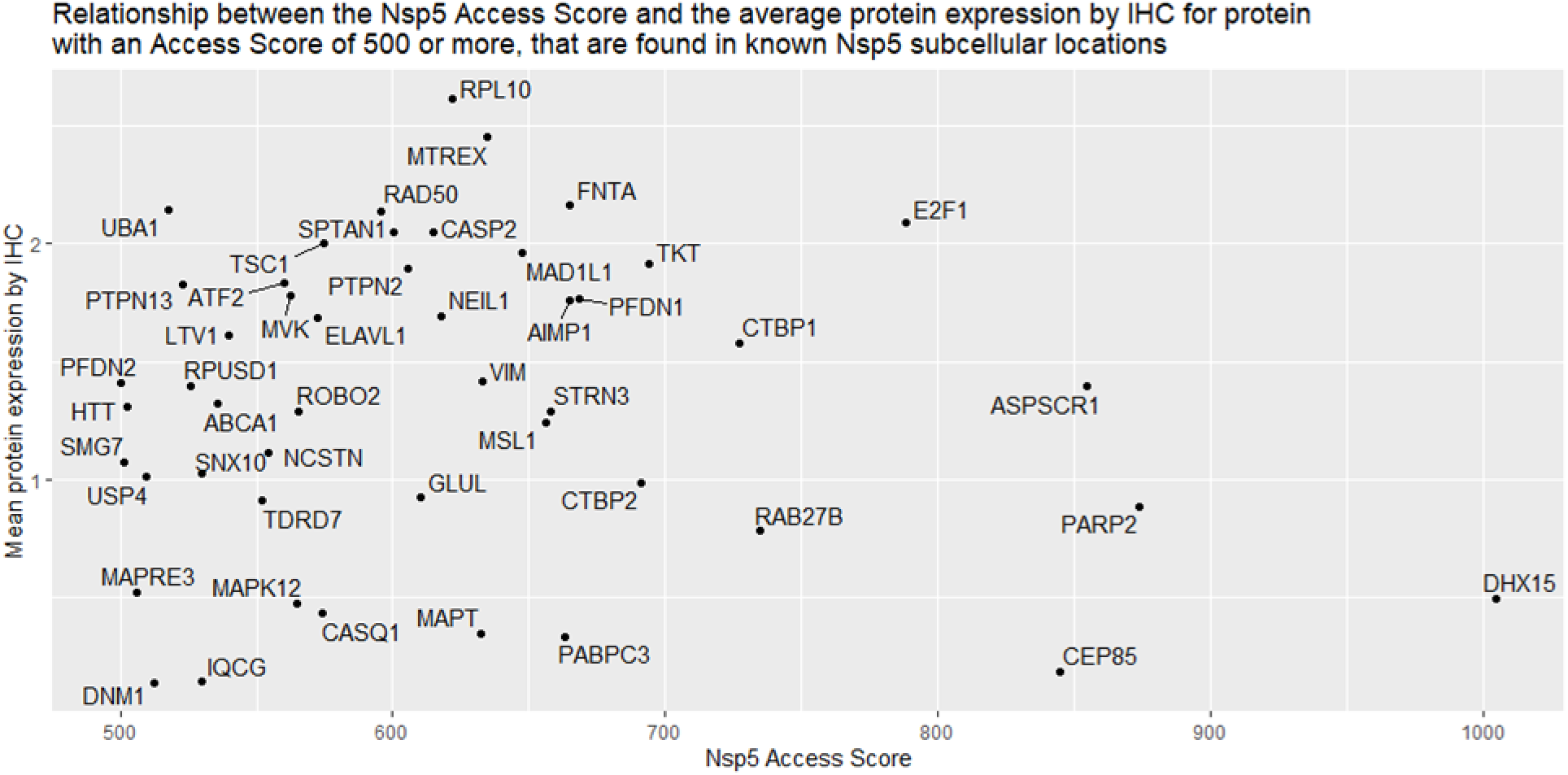
Proteins with a Nsp5 access score of 500 or more, that could be found in the same cellular compartment as Nsp5 (48 proteins), were plotted against their expression in the human body. For each protein, the mean expression by IHC is the mean across all tissues measured and reported in the HPA (Not detected = 0, Low = 1, Medium = 2, High = 3, Not measured = NA [which were ignored/removed]).

### Network Analysis and Pathways of Interest

Imputation in STRING of these 48 human proteins with a Nsp5 access score over 500 and plausible colocalization, revealed multiple pathways of interest (Fig. 6, Additional File 1: Table S15). The pathway containing the most proteins that may be targeted by and colocalize with Nsp5 was mRNA processing (DHX15, ELAVL1, LTV1, PABPC3, RPL10, RPUSD1, SKIV2L2, SMG7, TDRD7). Another prominent pathway was apoptosis, with multiple proteins involved directly in apoptosis or its regulation (CASP2, E2F1, FNTA, MAPT, PTPN13). DNA damage response, mediated through ATF2, NEIL1, PARP2, and RAD50 may also be targeted by Nsp5. PARP2 had the second highest Nsp5 access score in our analysis, and the predicted cleavage site at Q352 is located between the DNA-binding domain and the catalytic domain [73].

**Fig. 6.**
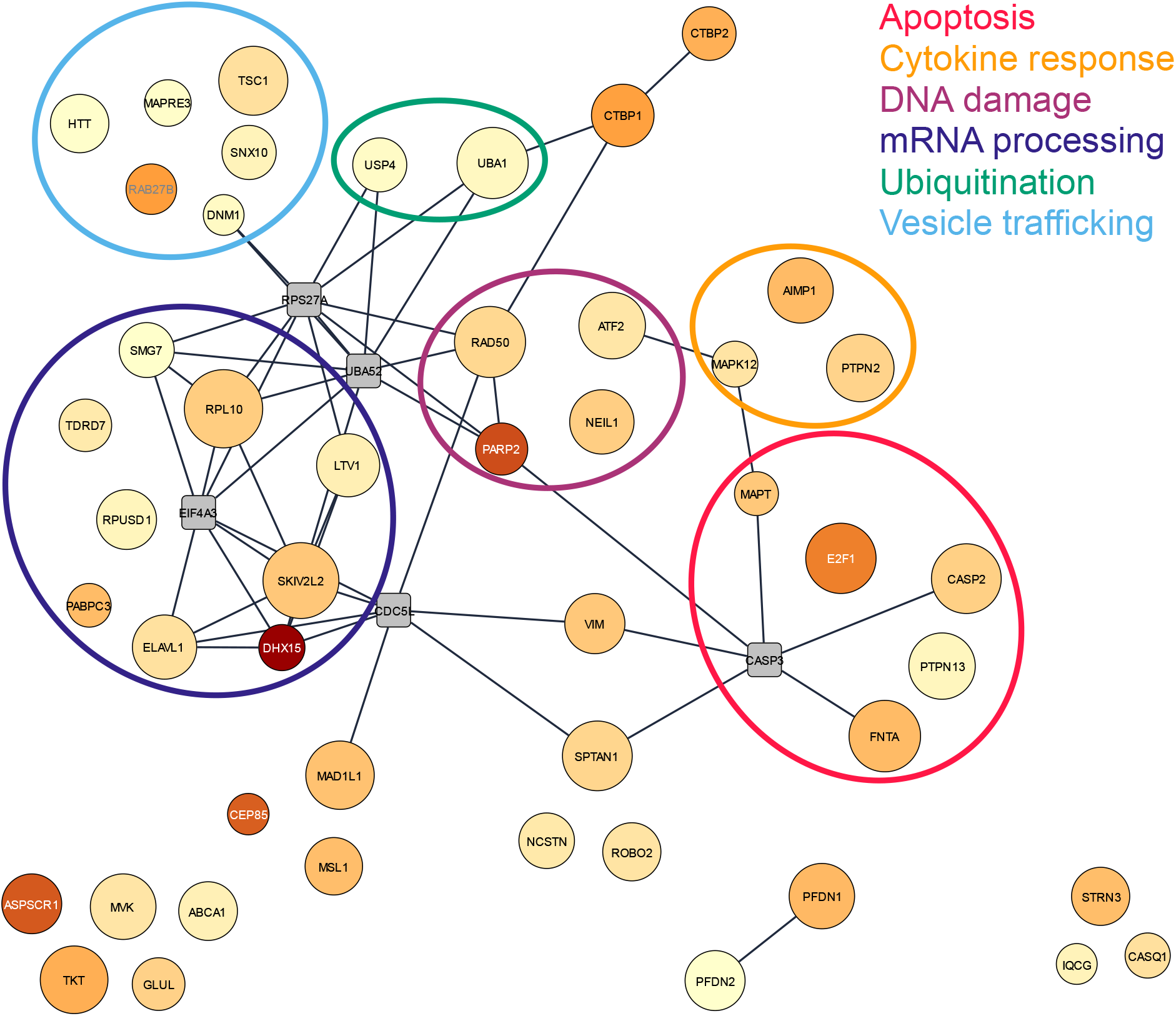
Network of proteins with plausible Nsp5 colocalization a Nsp5 access score above 500. Node color represents the Nsp5 access score (light yellow = 500, dark red = 1005). Node size indicates the mean expression across all tissue. Edge linking two nodes notes a known interaction between these proteins. Grey squares are proteins added by STRING to add connectivity to the network, but do not have an access score above 500 and/or plausible colocalization with Nsp5. Circles highlighting pathways were based on STRING gene set enrichment analysis coupled with manual searches in databases (Uniprot, GeneCARD, PubMed).

Proteins involved in membrane trafficking (RAB27B and SNX10), or in microtubule organization (DNM1, HTT, MAPRE3, TSC1) were also enriched in this focused dataset, which were grouped together under the descriptor “vesicle trafficking”. Two proteins related to ubiquitination (UBA1 and USP4) were also amongst these potential Nsp5 targets. Nsp3 mediated modulation of ubiquitination has been shown to be important for IFN antagonism [32–39], and there is also evidence for Nsp5 mediated reduction of ubiquitination [45, 47]. Finally, a group of proteins implicated in cytokine response was also strongly predicted to be cleaved (AIMP1, MAPK12, and PTPN2), which are involved in downstream signaling of multiple cytokines [74–77].

## Discussion

To provide context to the growing list of coronavirus-host protein-protein interactions, and to aid in the interpretation of experiments focused on human proteins cleaved by coronavirus Nsp5, we applied a bioinformatics approach to predict human proteins cleaved by Nsp5. Our proteome-wide investigation compliments *in vitro* experiments, which are limited to only a subset of potential human protein substrates based on what proteins are expressed by the cell type chosen, resulting in different proteins appearing to be cleaved by [47, 48], or interact with Nsp5 [49–54].

The NetCorona neural network generated long lists of potentially cleaved human proteins, but mismatches between these predictions and the *in vitro* mapping of Nsp5 cleavage sites indicated that NetCorona scores alone were insufficient for accurate predictions. Similar to previous reports [58–60], solvent accessibility helped to filter predictions based on primary sequence alone, which was made possible thanks to the PSAIA tool which automated the measurement of motif ASA with an easy-to-use GUI that handled batch input of PDB files [64].

Human proteins predicted to be cleaved by Nsp5 did not correlate with Nsp5-human protein-protein interactions identified *in vitro*, and Nsp5 overall appears to interact with fewer human proteins compared to other Nsps and structural proteins [51]. This may be because the proteolytic activity of Nsp5 reduces the efficiency of proximity labeling/affinity purification, whereby Nsp5 may cleave proteins it interacts with most favorably, reducing the appearance of host protein interactions. The small but statistically significant negative correlation between the strength of the Nsp5-human protein interaction and the human protein’s maximum NetCorona score may be evidence of this. Indeed, different sets of interacting proteins are obtained when using the catalytically inactive Nsp5 mutant C145A versus the wildtype Nsp5 [49, 51, 54]. We therefore hypothesize that the interactions observed by proximity labeling/affinity purification do not reflect Nsp5 mediated proteolysis and instead represent non-proteolytic protein-protein interactions, which may still be important to understanding Nsp5’s role in modulating host protein networks.

N-terminomics based approaches have identified many potential Nsp5 cleavage sites in human proteins [47, 48], but they have some limitations that bioinformatics can compliment. Trypsin is used in the preparation of samples for mass spectrometry, which generates cleavages at lysine and arginine residues that are not N-terminal to a proline. Lysine and arginine appear in many cleavage sites predicted by NetCorona, meaning that cleavage by trypsin may mask true cleavage sites by artificially generating a N-terminus proximal to a P1 glutamine residue. Only one protein overlaps between the Koudelka *et al*. and Meyer *et al*. results, as these studies used different cell lines, and thus different proteins will be expressed, and the methods of exposure to Nsp5 also differed (cell lysate incubated with Nsp5 vs SARS-CoV-2 infection of cells) [47, 48]. Meyer *et al*. point out that the lysate-based method used by Koudelka *et al*. strips proteins of their subcellular context, which may lead to observed cleavage events that are not possible *in vivo* during infection [48]. In contrast, our bioinformatics analysis is cell-type and methodology agnostic as it examined the entire human proteome. The cleavage sites predicted *in silico*, combined with knowledge of Nsp5 subcellular localization and protein networks, identified several interesting human proteins and pathways.

DHX15 contained a predicted cleavage site with the highest Nsp5 access score, and the protein may co-localize with Nsp5 in the nuclei of infected cells, making it a significant protein of interest. DHX15 is a DExD/H-box helicase, a family of proteins that serves to detect foreign RNA, triggering an antiviral response (reviewed by [78]). The role of DHX15 in anti-viral defense is diverse, including by binding viral RNA with NLRP6, which activates Type I/III interferons and IFN-stimulated genes in the intestine of mice [79]. DHX15 mediated sensing of viral RNA activates MAPK and NK-κB innate immune signaling [80], and also acts as a coreceptor of viral RNA with RIG-I that increases antiviral response and cytokine production [81].

DHX15 is not reported to be a significant interactor with SARS-CoV-2 proteins [51], however it is capable of binding both dsRNA and ssRNA [81], suggesting it could bind coronavirus ssRNA. The location of the predicted Nsp5 cleavage site (Q788) is very close to its C-terminus (Y795), so cleavage by Nsp5 would remove only seven amino acids. However, there is a SUMOylation site (K786) at P3 of the cleavage motif, and the de-SUMOylation of DHX15 results in increased antiviral signaling [82]. Therefore, it is possible that Nsp5 cleavage at Q788 may disrupt SUMOylation, by reducing the length of the peptide accessible by the SUMOylation complex, contributing to the general dysregulation of a coordinated innate immune response to viral infection.

PARP2 contained a predicted Nsp5 cleavage site with the second highest Nsp5 access score, at Q352. PARP2 is involved in DNA damage recognition and repair, and its correct functioning prevents apoptosis in the event of a double stranded break (DSB) [83]. PARP2 also plays a role in the adaptive immune system, as it helps prevent the accumulation of DSBs during TCRα gene rearrangements in thymocytes, promoting T-cell maturation [84]. The predicted Nsp5 site occurs between the DNA binding and catalytic domains of PARP2 [85], meaning the cleaved protein would be unable to recognize damaged DNA, contributing to apoptosis. Interestingly, this is similar to the native activity of human caspase-8, which cleaves PARP2 in its DNA binding domain during apoptosis [86]. SARS-CoV-2 infection of lung epithelial cells was found to increase caspase-8 activity, resulting in cleaved PARP1, a homolog of PARP2 [87]. SARS-CoV Nsp5 activity is known to be pro-apoptotic, via the activation of caspase-3 and caspase-9 [88]. Overall, if coronavirus Nsp5 cleaves PARP2, it could contribute to the pro-apoptotic cell state observed in infected cells.

Proteins with roles in cytokine response and antiviral defense were also identified as strongly predicted targets of Nsp5 cleavage (AIMP1, MAPK12, PTPN2). AIMP1 is crucial to antiviral defense in mice [74], and it is targeted for degradation by hepatitis C virus envelope protein E2 [75]. MAPK12 function appears to be important for SARS-CoV-2 replication, as knockdown by siRNA results in lower virus titers [76]. This phenotype is similar to the knockdown of Nsp5 targeted proteins resulting in lower virus replication, as observed by Meyer *et al*. [48]. T-cell protein tyrosine phosphatase, PTPN2, negatively regulates the antiviral response of MITA [89] and the JAK-STAT pathway of the innate immune system [90]. Knockout of PTPN2 results in systemic inflammatory responses in mice resulting in premature death [91], and a genetic polymorphism resulting in PTPN2 loss of function increases ACE2 expression, resulting in greater susceptibility to SARS-CoV-2 infection [77].

The modulation of mRNA processing, DNA damage recognition/apoptosis, and cytokine signaling stood out as the most interesting pathways that may be influenced by Nsp5 cleavage. As coronaviruses are known to bias mRNA processing, trigger apoptosis, and disrupt innate immune responses (reviewed by [15]), these results suggest that Nsp5 mediated cleavage may aid in the molecular pathogenesis of diseases caused by coronaviruses diseases. These pathways, and specific proteins mentioned above, represent interesting targets for further analysis *in vitro*.

## Limitations

Recent studies comparing Nsp5 proteases from various coronaviruses have indicated that despite sharing significant sequence and structural similarity, they cleave and interact with different human proteins *in vitro* [47, 54]. In general, a pan-coronavirus predictor of Nsp5 cleavage sites may not be feasible. For example, SARS-CoV-2 Nsp5 accommodates more diversity at P2 than SARS-CoV Nsp5 [25], which would influence the human proteins that could be cleaved. Refinements to the NetCorona neural network to improve its predictive accuracy, or make them virus-specific, would be beneficial and have recently been attempted [92]. Nsp5 also likely accommodates more diversity at P1’ and P3 than what NetCorona was trained on, based on the naturally occurring 1ab variants we report here, and the human proteins observed to be cleaved *in vitro*. That histidine can be accommodated at P1 and phenylalanine at P2, albeit unfavorably, further adds complexity to what human proteins may be cleaved by Nsp5 [17, 25, 47, 61]. The results of this study are instead meant to provide guidance for *in vitro* experimental design and interpretation of experimental results, in addition to suggesting proteome-wide trends in molecular pathways that Nsp5 mediated cleavage may disrupt.

## Conclusions

The large volume of recent coronavirus research and data requires proteome-wide views of interpretation. The results of this study are intended to compliment the various *in vitro* approaches that have been used to identify Nsp5-human protein interactions, and to map specific Nsp5 cleavage sites in human proteins. As Meyer *et al*. discuss, specific targeting of proteins by Nsp5 appears likely, as the knockdown of certain Nsp5-targeted proteins reduces viral reproduction [48]. We have built upon the original NetCorona study by performing detailed structural analysis of predicted cleavage sites, and protein network and pathway analysis. Coronavirus Nsp5 was predicted to play a role in the targeted disruption of mRNA processing, cytokine response, and apoptosis, which are interesting targets for future analysis and characterization. We hope that our analysis and the proteome-wide datasets generated will aid in the interpretation and design of additional experiments towards understanding Nsp5’s role in coronavirus molecular pathology.

## Methods

### Amino Acid Sequence Datasets

Three datasets of amino acid sequences of human proteins were downloaded from the UniProt reference human proteome on April 14 2020: “All Human Proteins” contains 74811 human protein sequences including splice variants and predicted proteins; “One Protein Per Gene” contains 20595 human protein sequences where only one sequence is provided for each gene; “Proteins With PDB” is the “All Human Proteins” dataset filtered in UniProt for associated entries in the RCSB Protein Data Bank, generating a dataset of 6806 human protein sequences. 9404 amino acid sequences of the SARS-CoV-2 1ab polyprotein were obtained from NCBI Virus on August 4 2020. Sequences with inconclusive “X” residues were filtered out as they were not correctly handled by NetCorona, leaving 8017 SARS-CoV-2 1ab polyprotein sequences to be analyzed.

### NetCorona Analysis

The command line version of NetCorona was used to predict Nsp5 cleavage site scores for human and viral protein sequences [55]. To overcome the input file limit of 50,000 amino acids per submission and handle sequences with non-standard amino acids, a Python script was developed. This script partitions the input data, runs the NetCorona neural network on each subset, and parses and concatenates the output data. The output file includes sequence accession number, position of P1-Glutamine (Q) residue, Netcorona score (0.000-1.000) and a 10 amino acid sequence motif of positions P5-P5’ (Additional File 2-4). Note that the neural network itself uses positions P5-P4’ (9 residues) for calculating the score. Gene names and other identifiers associated with each UniProt ID containing a NetCorona score >0.5 were collected in Microsoft Excel (Additional File 1: Table S2-S4). Scores from NetCorona run on each dataset of proteins were parsed and compared with a Kolmogorov-Smirnov (KS) test, to assess the null hypothesis that the scores for each population are drawn from the same distribution. Unique glutamine motifs from 1ab polyprotein sequences were identified using Microsoft Excel (Additional File 1: Table S1). Statistical analysis and the generation of graphs was performed using GraphPad Prism (version 9.1.0)

### Structural Analysis

PDB metadata associated with proteins in the “Proteins With PDB” dataset that also contained a predicted Nsp5 cleavage (NetCorona score >0.5), were downloaded from the RCSB PDB website by generating a custom report in .csv format. Homology models, and structures with a resolution greater than 8Å or where resolution was not reported, were removed. Nsp5 cleavage sites predicted by NetCorona were matched with one PDB file per cleavage site, by searching the PDB metadata for the predicted 9 amino acid cleavage motif using Microsoft Excel (Additional File 6). The entire predicted 9 amino acid motif must appear in the PDB file to be considered a match. Matches between a PDB file and predicted cleavage motif were manually corrected when the motif sequence appeared by chance in a PDB containing the incorrect protein.

PDB files containing a predicted Nsp5 cleavage site were then batch downloaded from the RCSB PDB, and analyzed 100 at a time using the Protein Structure and Interaction Analyzer (PSAIA) tool using default settings [64], with chains in each PDB analyzed independently. The total accessible surface area (ASA) of each residue was calculated using a Z slice of 0.25 Å and a probe radius of 1.4 Å. XML files output by PSAIA were combined in Microsoft Excel, to create searchable datasets for each 9 amino acid motif predicted to be cleaved by NetCorona, and the total ASA of all atoms in each 9 amino acid motif were summed. The motif’s ASA was then multiplied by the NetCorona score to provide a Nsp5 access score.

Proteins known to be cleaved by mammalian chymotrypsin-like proteases were independently obtained from the RCSB PDB, and the known cleaved motifs were analyzed as above. Protein structures and homology models of SARS-CoV-2 proteins were obtained the RCSB PDB and from SWISS-MODEL [93] and were analyzed as above. Publication quality figures were generated using PyMOL 2.3.0.

### Tissue Expression and Subcellular Localization Analysis

Proteins with Nsp5 access score above 500 were loaded into the STRING app [65] within Cytoscape [66] (version 1.6.0 and 3.8.2 respectively) using Uniprot ID, Homo sapiens background, 0.80 confidence score cut-off and no additional interactor for the pathway enrichment analysis (Additional File 1: Table S13-S15). The node table (including tissue expression scores and compartments score for each protein) was exported to R for wrangling and data visualization using the tidyverse and ggrepel packages [94–96].

To increase confidence, tissue expression and subcellular localization data were obtained from the Human Protein Atlas which are all based on immunohistochemistry (tissue expression) or confocal microscopy (subcellular localization) [72, 97]. Each entry was then matched in R, table joining was done using Uniprot IDs. Expression levels noted as “Not detected”, “Low”, “Medium” or “High” were replaced by numeric values ranging from 0 to 3. Mean expression was calculated as the mean expression across all tissues, removing missing values from the analysis.

The following intracellular locations were used to encompass the nucleus, cytoplasm and endoplasmic reticulum: “Cytosol”, “Nucleoplasm”, “Endoplasmic reticulum”, “Microtubules”, “Nuclear speckles”, “Intermediate filaments”, “Nucleoli”, “Nuclear bodies”. All the proteins that did not include one or more of these locations in the HPA database were excluded from further analysis.

### Protein Network Analysis

The 48 proteins with a Nsp5 access score >500 and that had the potential to be found in the same cellular compartment as Nsp5 were imported into the STRING app (again within Cytoscape) while allowing a maximum of 5 additional interactor for the network generation instead of none. All the other parameters were left unchanged. Individual nodes that had no protein-protein interactions with other proteins in the network were manually moved closer to other nodes presenting the same or similar pathway. When proteins could interact in multiple pathways represented here, a “main pathway” was assigned based on literature search. Node color was a gradient based on Nsp5 access score. Node size increased with the mean expression. Edges represent protein-protein interaction (confidence > 0.80). Gene name labels were colored based on the Nsp5 access acore for readability only.

## Acknowledgements

We wish to thank the Crowdfight COVID (crowdfightcovid19.org) for facilitating this collaboration, and Ian M. Boyce for assistance editing figures. The National Institute of Standards and Technology (NIST) notes that certain software and materials are identified in this article to explain a procedure as completely as possible. This does not imply a recommendation or endorsement by NIST, nor does it imply that the particular materials are necessarily the best available for the purpose. The opinions expressed in this article are the authors’ own and do not necessarily represent the views of NIST.

## Author Contributions

BMS conceived the idea and supervised the study. BMS, VL, DGB, NSB generated the data. BMS, VL, PDT analyzed the data. BMS, VL, DGB generated the figures. BMS and VL drafted the manuscript. All authors edited the manuscript. BMS acted as corresponding author. All the authors have read and approved the final manuscript.

## Availability of Data and Materials

PSAIA was obtained here: http://complex.zesoi.fer.hr/index.php/en/10-category-en-gb/tools-en/19-psaia-en

## Additional Files

Additional files are available here: https://doi.org/10.6084/m9.figshare.14751306

**Additional File 1:** This includes Supplementary Tables S1-S15

**Additional File 2:** human_all_proteins_netcorona.txt (108MB)

This is the raw data output by NetCorona following analysis of the “All Human Proteins” dataset

**Additional File 3:** human_one_gene_netcorona.txt (46MB)

This is the raw data output by NetCorona following analysis of the “One Protein Per Gene” dataset

**Additional File 4:** dataset human_w_PDB_netcorona.txt (16MB)

This is the raw data output by NetCorona following analysis of the “Proteins With PDB” dataset

**Additional File 5: Figure S1** interaction scores vs max NetCorona score.pdf

Nsp5-human protein interaction data from Samavarchi-Tehrani *et al*. [51], plotted against the maximum NetCorona score for human proteins from the “One Protein Per Gene” dataset.

**Additional File 6:** matching predicted motif to PDB.xlsx

Raw data displaying how predicted cleaved motifs were matched to a PDB file

**Additional File 7: Figure S2** ASA vs NetCorona score.pdf

Accessible surface area (ASA) of a predicted and known Nsp5 motifs plotted against NetCorona scores, with data published by Moustaqil *et al*. and Koudelka *et al*. highlighted [46, 47], and the Nsp5 access score cut-off displayed.

